# Pre-treatment Microbiome Diversity and Function is associated with Expansion of Cytotoxic and Regulatory Immune Populations after N-803 treatment in People with HIV

**DOI:** 10.1101/2025.10.01.679827

**Authors:** Ashma Chakrawarti, Ross T. Cromarty, Christopher M. Basting, Jodi Anderson, Ty A. Schroeder, Kevin Escandon, Robin Shields-Cutler, Robert Langat, Erik Swanson, Patrick Soon-Shiong, Jeffrey T. Safrit, Leonard S. Sender, Sandeep Reddy, Jeffrey S. Miller, Joshua Rhein, Timothy W. Schacker, Nichole R. Klatt

**Affiliations:** Department of Surgery, Division of Surgical Outcomes and Precision Medicine Research, University of Minnesota, Minneapolis, MN; Department of Medicine, University of Minnesota, Minneapolis, MN; Department of Biology, Macalester College, Saint Paul, MN; ImmunityBio, Culver City, CA, USA

**Keywords:** N-803, Fecal microbiome, Baseline diversity, SCFAs, Immune activation markers, Immunotherapy, *Faecalibacterium*, HIV reservoirs, cure therapy

## Abstract

**Background:** N-803, an IL-15 superagonist, is currently being studied in clinical trials as a treatment to reverse HIV latency. However, its effects on the gut microbiome are not well understood.

**Methods:** In this longitudinal metagenomic study, we analyzed fecal microbiomes from ART-suppressed people with HIV at four different timepoints before, during, and after N-803 treatment.

**Results:** Overall taxonomic and functional diversity did not change significantly, yet beneficial microbial taxa and pathways were enriched after N-803. Specifically, the relative abundance of *Faecalibacterium prausnitzii* increased significantly after N-803, whereas histidine degradation pathways, often associated with pro-inflammatory mucosal state, decreased. A higher baseline microbial diversity correlated with stronger CD8^+^ and natural killer (NK) cells activation and reduced frequency of rectal HIV RNA^+^ cells. *MaAsLin2* analyses further associated short-chain fatty acid (SCFA)-producing taxa and pathways with increased immune activation markers.

**Conclusions:** These results indicate that gut microbiome diversity prior to immunotherapy influences host response and suggest that microbiome-based strategies could improve efforts to cure HIV.

## Background

Antiretroviral therapy (ART) has transformed Human Immune Deficiency Virus (HIV-1) infection into a chronic, manageable condition. However, ART alone cannot eradicate latent viral reservoirs that persist primarily in CD4^+^ T cells within secondary lymphoid tissues and gut-associated lymphoid tissues (GALT) [1–3]. Even during long-term ART, these reservoirs remain active and produce low-level viral transcripts, which contribute to ongoing immune activation and inflammation in people living with HIV (PWH) [4,5]. Further, this contributes to cardiovascular disease, metabolic disorders, neurocognitive impairment, and accelerated aging [6].

Immunotherapies that reduce the HIV reservoir and boost natural killer (NK) and CD8+ T cell functions are of interest, given that these immune functions are impaired in PWH [7–9]. N-803 (ImmunityBio, USA), an IL-15 superagonist, is designed to stimulate NK and CD8+ T cell proliferation and activation [10–12]. Originally developed for cancer therapy, N-803 has shown promise in preclinical SIV studies and early HIV trials [13–15] by enhancing antiviral effector function and inducing viral protein expression in latently infected cells [15–17]. In our recent Phase 1B trial, three subcutaneous doses of N-803 enhanced NK cell numbers and function, which associated with decreased viral RNA+ CD4+ T cells in lymphoid and gut tissues [18].

Findings from oncology research have shown that the gut microbiome can influence host response to immune-based treatments [19–21]. These effects are thought to be mediated by the production of microbial metabolites (e.g., short-chain fatty acids, SCFAs), antigenic mimicry between microbial and host antigens, and regulation of immune cell subsets [20,22,23]. SCFAs, particularly acetate, propionate, and butyrate, produced by gut commensals including *Faecalibacterium prausnitzii, Roseburia*, and members of the *Lachnospiraceae* family, are reported to enhance epithelial barrier integrity, reduce inflammation, and modulate mucosal immunity [21,24,25]. In HIV infection, both microbial dysbiosis and translocation are well-documented contributors to immune dysfunction and chronic inflammation [26,27], even during effective ART [28,29]. However, whether these factors influence immunotherapy outcomes in chronic HIV is poorly understood.

To address this knowledge gap, we conducted a longitudinal shotgun metagenomic study of microbiome dynamics following N-803 treatment. By integrating microbial diversity, functional pathway analysis, immunological profiles, and virological measurements, we aimed to determine whether microbiome-immune interactions influence responses to immunotherapy and whether N-803 itself induces compositional or functional shifts in the microbiome. Our analyses revealed a significant correlation between baseline gut microbial diversity, enhanced immune activation, and lowered frequency of HIV RNA^+^ cells, highlighting the microbiome as a potential modulator of therapeutic responses in ART-suppressed PWH receiving N-803.

## Methods

### Study Design and Participants

This study used fecal samples collected from participants in a Phase 1B clinical trial that evaluated the immunomodulatory effects of N-803, an IL-15 superagonist, in PWH (ClinicalTrials.gov Identifier: NCT04808908) [18]. The clinical trial enrolled adult participants who had stable ART with consistent viral suppression and maintained CD4+ T-cell counts above 350 cells/μL. All enrolled individuals received three doses of subcutaneous N-803 (6 μg/kg) administered at 21-day intervals. The clinical trial protocol was approved by the University of Minnesota Institutional Review Board (IRB) (UMN IRB STUDY0007810), and all participants provided informed consent. Detailed clinical inclusion and exclusion criteria, as well as primary clinical outcomes, are fully described in the clinical trial paper [18].

### Fecal Sample Collection and Shotgun Metagenomics

Fecal samples for microbiome analysis were longitudinally collected from participants at defined intervals throughout N-803 administration (Figure 1). Samples were collected before the initial dose of N-803 (*Baseline)*, following the second dose (*Dose2*), immediately after the completion of all three doses (*Postdrug*), and at a follow-up visit approximately three months post-treatment (*Month3*). Shotgun metagenomic sequencing was performed on an Illumina NovaSeq SP platform by the University of Minnesota Genomics Core (UMGC). Taxonomic profiles were generated using MetaPhlAn4, and microbial pathways were annotated via HUMAnN3. Full sequencing and preprocessing protocols are detailed in Supplementary Materials.

**Figure 1.**
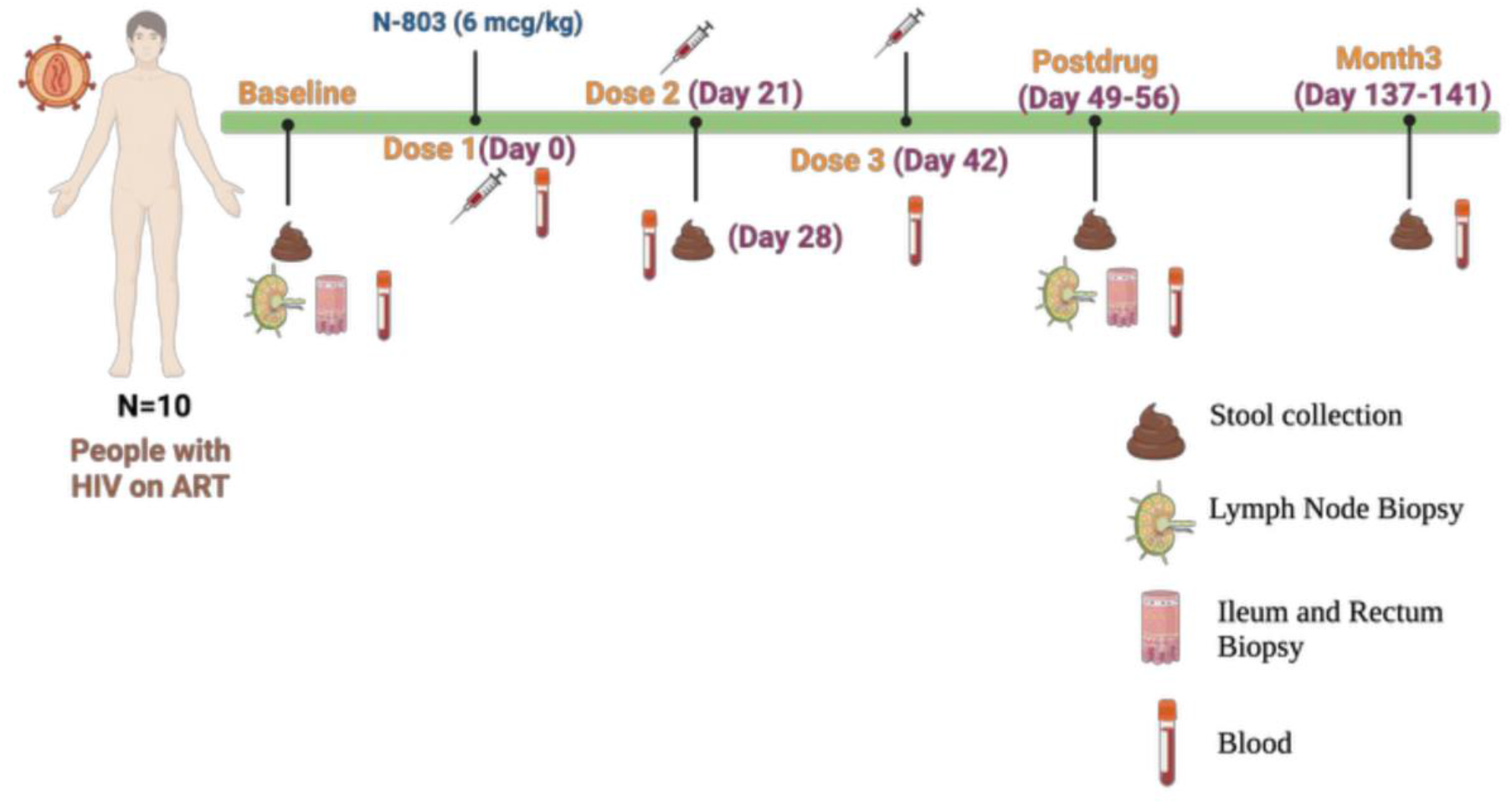
Schematic overview of N-803 administration schedule and longitudinal fecal sample collection for gut microbiome profiling. Participants received three subcutaneous doses of N-803 (6 μg/kg) at 3-week intervals. Stool samples were collected at four timepoints: Baseline (pre-treatment), Dose2 (~Day 28), Postdrug (~Day 49-56), and Month3 follow-up (~Day 137-141). Created in BioRender. Chakrawarti, A. (2025) https://BioRender.com/5u80knn

### Additional Clinical Methods

The frequency of HIV RNA^+^ (vRNA^+^) cells was quantified in lymph node, ileum, and rectum biopsies using RNAScope in situ hybridization. Positive cells were normalized to tissue mass and expressed as vRNA^+^ cells per gram of tissue. Immune profiling using mass cytometry by time of flight (CyTOF) was performed as part of the clinical trial study. These methods, including detailed protocols, are fully described in the clinical trial paper [18].

### Statistical Analysis

Alpha diversity was quantified using Shannon and Simpson indices, calculated on both taxonomic and functional abundances, and temporal trends were assessed using linear mixed-effects models with participants as a random intercept. Overall effects of sampling time (Baseline, Dose2, Postdrug, Month3) were evaluated by ANOVA and pairwise comparisons between timepoints with Benjamini-Hochberg (BH) adjustment for multiple comparisons. Beta diversity was calculated using Aitchison distances on CLR-transformed abundance tables and Bray-Curtis dissimilarities on proportional abundance data. Compositional shifts were visualized using Principal Coordinates Analysis (PCoA), and statistical significance across timepoints was evaluated by pairwise PERMANOVA tests with BH-adjustment. Differential abundance analysis was performed using mixed-effects linear modeling (*MaAsLin2*), controlling for participant variability as a random effect, and BH-adjusted q-values ≤0.25 were considered significant.

To assess the relationship between baseline microbiome diversity and post N-803 immune responses, we performed Spearman correlation analyses between Baseline Shannon diversity and i) the Postdrug expression levels of immune markers on NK cells and CD8+ T cells, and ii) the percent change in marker expression from Baseline to Postdrug for each participant. These included markers associated with cytotoxicity (Perforin, Granzyme B), activation (CD38, HLA-DR), proliferation (Ki-67), differentiation and senescence (CD27, CD57), inhibitory signaling (PD-1, NKG2A), and apoptosis (FasL). For correlation analyses, nominal p-values <0.05 were considered significant.

BH-adjusted q-values ≤0.25 were considered significant for all analyses with multiple-hypothesis correction due to the small sample size. All analyses were performed using R (v4.3.3). Full analytical details, including software packages, parameters, and data visualization methods, are provided in the Supplementary Materials.

## Results

### Study Cohort and Sample Collection

Fecal metagenomic data were generated from ten virologically-suppressed PWH who completed all three subcutaneous doses of N-803 in a Phase 1B trial. Eight out of ten participants collected stool at every timepoint, with two participants missing the Month3 follow-up collection. Comprehensive immunological parameters and clinical outcomes are reported in the clinical trial paper [18].

### N-803 does not perturb the overall taxonomic structure and function of the gut microbiome

We generated shotgun metagenomic data from 38 stool samples to evaluate the effect of N-803 administration on fecal microbiome composition across four timepoints: Baseline, Dose2, Postdrug, and Month3. Taxonomic profiling demonstrated that the relative abundance of the top 20 bacterial genera remained largely stable over the study period, with no consistent treatment-related shifts observed (Figure 2A). The genus-level microbiota was dominated by *Bacteroides, Prevotella, Segatella, Roseburia*, and *Faecalibacterium*. Two participants lacked the genus *Segatella* or *Prevotella* among their top 20 genera and instead exhibited a higher relative abundance of *Bacteroides* and *Bifidobacterium* compared to the rest of the cohort.

**Figure 2:**
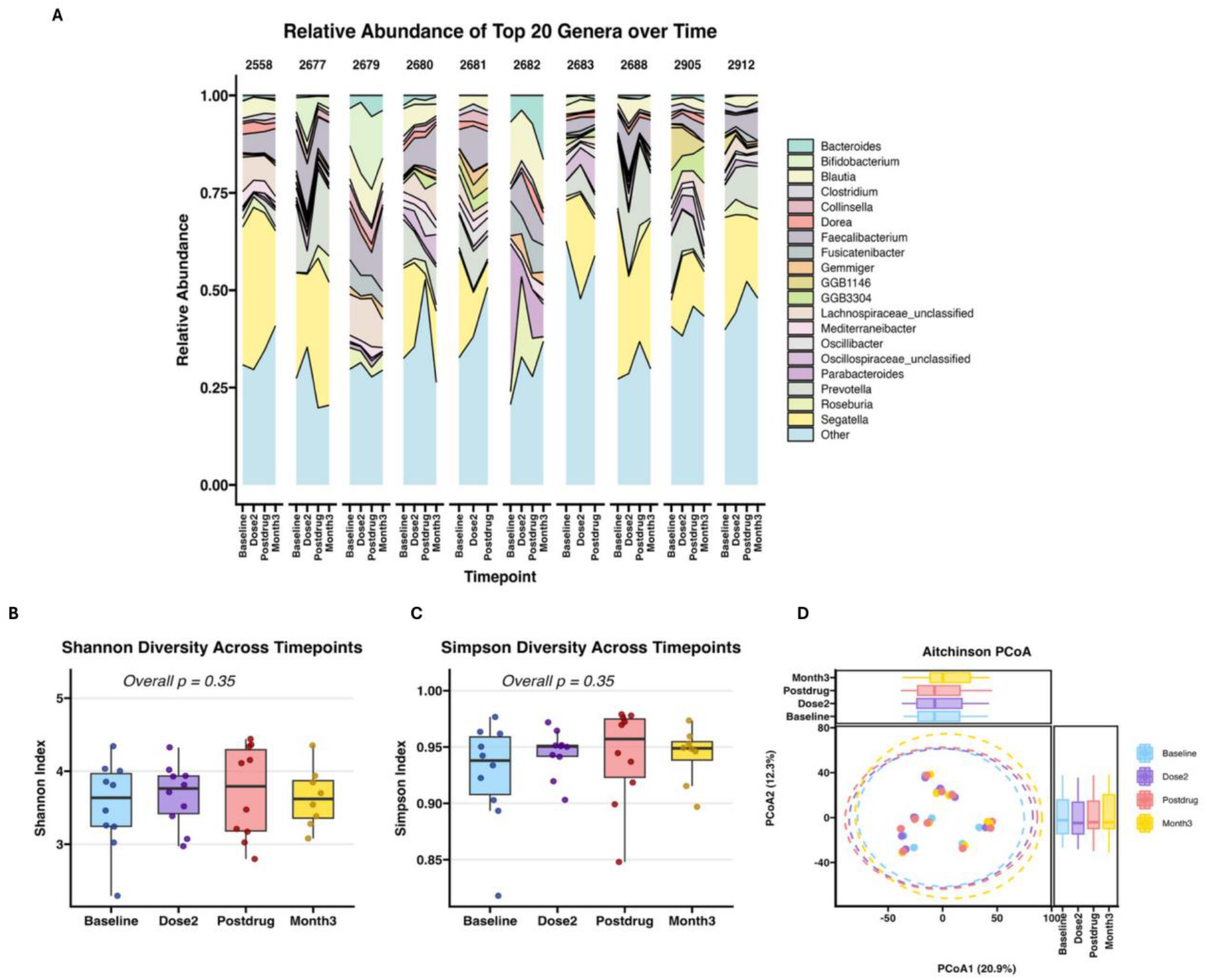
N-803 treatment on fecal microbiome composition and diversity. **(A)** Stacked area plot showing on the y-axis relative abundance (%) of bacterial genera detected in fecal samples from each participant across four sampling timepoints (Baseline, Dose2, Postdrug, and Month3 follow-up) on the x-axis. Genera represented in the legend include only the 20 most abundant taxa. **(B-C)** Boxplots of Shannon diversity **(B)** and Simpson index **(C)** were calculated from species-level relative abundance data over time. Overall, p-values indicate the fixed effect of timepoint. **(D)** Principal coordinates analysis (PCoA) of CLR-transformed species-level abundance data using Euclidean distances. Points represent samples colored by timepoint, with 95% confidence ellipses shown. Side boxplots display the distribution of PCoA scores for each timepoint along the first (x-axis, top) and second (y-axis, right) principal coordinate axes, highlighting within-group variability and overlap in community structure. No significant pairwise differences in community composition were detected between timepoints (pairwise PERMANOVA, *q* >*1* for all comparisons).

Shannon and Simpson indices were calculated to assess alpha diversity across the four timepoints. For both Shannon and Simpson diversity, the linear mixed-effects model indicated a non-significant overall effect of timepoint (*p* > *0*.*05*, ANOVA, Figure 2B-C). Pairwise comparisons with BH correction also had no statistically significant differences between any timepoints for either metric (*q* > *0*.*25*, see Supplementary Materials Figure S2 A-B). Similarly, beta diversity analyses showed no distinct clustering of microbial communities by timepoint, though each patient’s timepoint samples generally clustered together (Figure 2D). Statistical comparisons using pairwise PERMANOVA confirmed no significant compositional differences among any of the timepoints (*q* =1.0). Overall, these results suggest that N-803 administration had no detectable impact on the composition of the participants’ gut microbiota.

To evaluate whether N-803 influences the metabolic pathways present in the fecal microbiome, we calculated alpha diversity indices for MetaCyc pathway abundance derived from HUMAnN3 outputs. The Shannon and Simpson diversity metrics demonstrated that pathway abundances were stable across the four treatment timepoints, with no significant differences found in pathway-level richness or evenness (Shannon index, *p* = 0.69; Simpson index, *p* = 0.87) (see Supplementary Materials Figure S2 C-D). All pairwise comparisons among timepoints yielded non-significant results (*q* > 0.25). Likewise, beta diversity ordination of the metabolic pathway profiles did not demonstrate distinct clustering by timepoint (see Supplementary Materials Figure S3); pairwise PERMANOVA comparisons also indicated no significant compositional differences (*q ≥ 0*.*99*).

### Taxonomic and microbial pathway shifts following N-803 administration

Linear mixed effects modeling using *MaAsLin2* identified two species whose abundances changed after N-803 administration (Figure 3A). The relative abundance of *Segatella sinensis* significantly declined by Month3 (*q* = 0.068). In contrast, *Faecalibacterium prausnitzii* increased by Dose2 (*q* = 0.225) and remained elevated at Month3 (*q* = 0.129).

**Figure 3:**
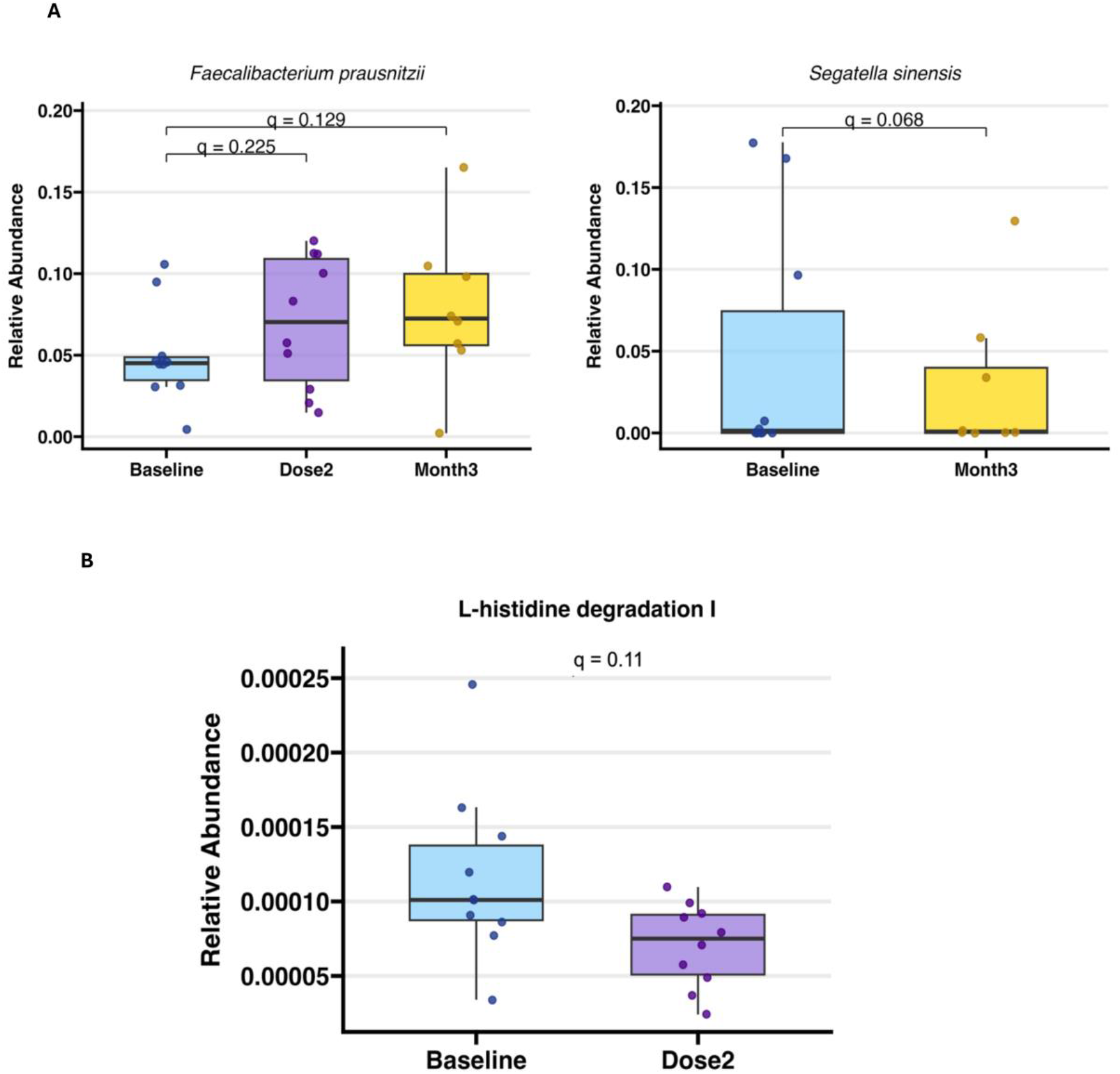
Differential abundance of fecal bacterial taxa and MetaCyc pathway after N-803 therapy. Box-and-whisker plots display the relative abundances returned by **(A)** MetaPhlAn 4 (taxa) and **(B)** HUMAnN 3 (pathway). They satisfied the significance threshold (*q* < *0*.*25*). Baseline samples are plotted in blue and post-drug samples in purple (Dose2) or yellow (Month3).

At the functional pathway level, the relative abundance of microbial histidine degradation decreased by 30% from baseline after N-803 (*q* = 0.11, Figure 3B). In contrast, SCFA pathways, which play roles in supporting the mucosal barrier and regulating the immune system, showed modest increases, although these changes did not reach significance after FDR adjustment (nominal *p* < 0.05, *q* > 0.25; Supplementary Materials Figure S4, Supplementary Materials Table S2).

### Fecal microbiome diversity at baseline correlates with N-803 treatment outcome

We next tested whether baseline fecal microbial diversity modulates immune responses to N-803 treatment. Correlation analyses compared Baseline Shannon diversity with Postdrug CD8^+^ T cell and NK cell frequencies and functional expression markers. Baseline Shannon diversity showed a modest, non-significant positive trend (ρ = 0.267, *p* = 0.455) with Postdrug CD8^+^ T cell frequency (Fig. S5A). However, Baseline diversity significantly correlated with Postdrug CD69 expression level (ρ = 0.66, *p* = 0.0186; Figure 4A). Other markers, including NKG2A, CD16, and Ki-67, showed positive but non-significant trends (Fig. 4A).

**Figure 4:**
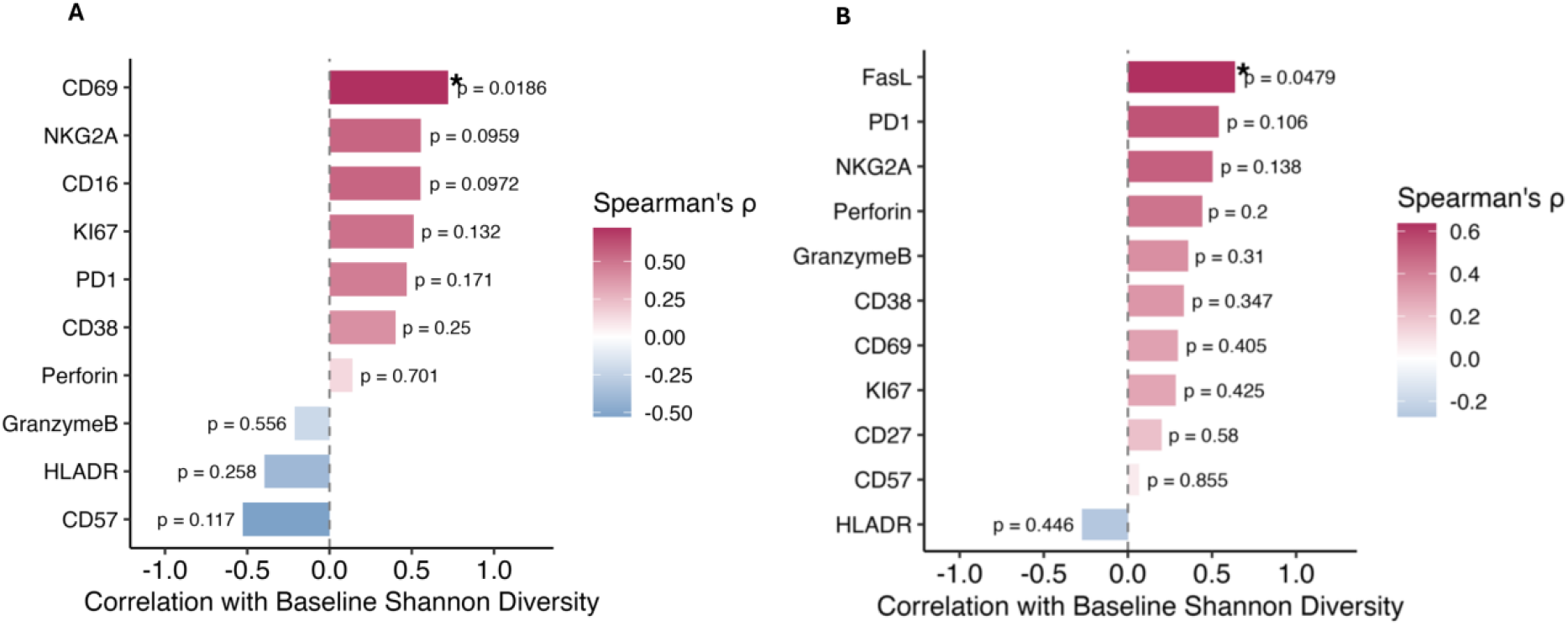
Baseline Fecal Microbiome Diversity Correlates with Post N-803 A) CD8^+^ B) NK cells Functional Markers. Barplot of Spearman’s correlation coefficients (ρ) between Baseline gut microbiome alpha diversity (Shannon diversity index) and post-treatment expression levels of immune activation markers measured by CyTOF. Bars to the right (positive ρ) increased immune activation; bars to the left (negative ρ) indicate inverse associations. Spearman correlation p-values are shown beside each bar; * indicates *p* < 0.05.

Similar findings were observed for NK cells, where Baseline Shannon diversity positively correlated with the Postdrug expression of FasL (ρ = 0.61, *p* = 0.0479). Other NK cell markers, including PD-1, NKG2A, and Perforin, displayed positive yet statistically non-significant trends, whereas Postdrug NK cell frequencies (see Supplementary Materials Figure S5A) and the expression of activation markers CD38 and CD69 showed no discernible correlation with baseline Shannon diversity. To further evaluate the hypothesis that Baseline microbial diversity influences the magnitude of immune modulation after N-803, we assessed the percent change in immune-marker expression from Baseline to Postdrug. We observed that Baseline Shannon diversity correlated with the percent change in CD8^+^NKG2A^+^ T cells (ρ = 0.89, *p* = 0.033) (Fig. S5A). This suggests that PWH harboring greater baseline microbial community diversity may exhibit more robust regulatory T-cell responses after N-803 treatment.

### Microbial diversity and specific microbial taxa are associated with reductions in vRNA^+^ cells in tissue from PWH

Given these observed microbiome-immune interactions, we next examined whether baseline gut microbial diversity similarly impacted virological outcomes, specifically tissue-associated viral transcription. We assessed correlations between Baseline Shannon diversity and tissue-associated frequency of vRNA^+^ cells in the rectum, ileum, and lymph nodes. Postdrug vRNA+ cell frequency inversely correlated with alpha diversity across all three tissue sites, with the strongest correlations observed in the ileum (ρ = −0.48, *p* = 0.162) and lymph nodes (ρ = −0.46, *p* = 0.187), though these trends did not reach statistical significance (see Supplementary Materials Figure S6 C-E). However, Baseline Shannon diversity was significantly correlated with the percent reduction in frequency of vRNA+ cells in the rectum (ρ = −0.67, *p* = 0.039) (Figure 5B), with participants who had higher Baseline microbiome diversity showing greater decreases in frequency of vRNA^+^ cells in the rectal mucosa. Negative trends were observed as well for the ileum (ρ = −0.21, p = 0.56) and lymph nodes (ρ = −0.60, p = 0.242; see Supplementary Materials Figure S6A-B).

**Figure 5:**
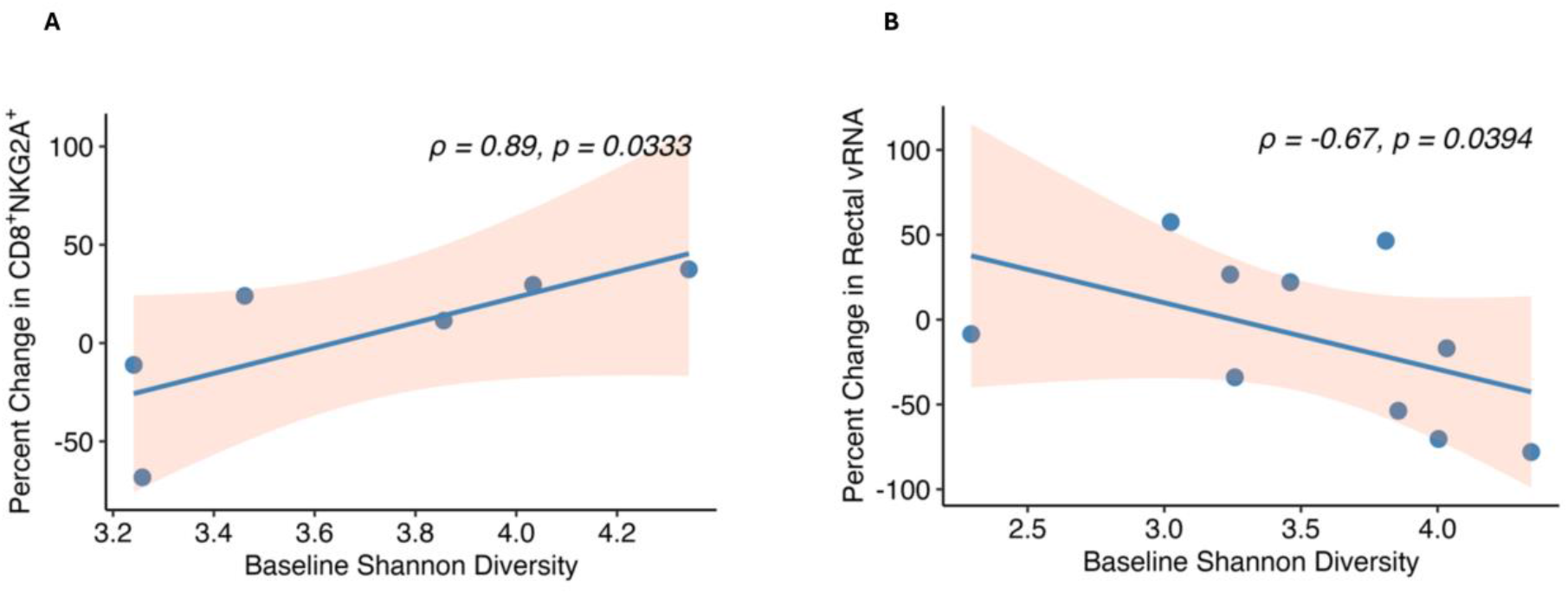
Baseline Fecal Microbiome Diversity Correlates with Post N-803 (A) Percent change in CD8+NKG2A+ (B) Percent change in frequency vRNA^+^ cells of rectum. The left panel (**A**) shows the positive correlation between Baseline Shannon diversity and percent change in CD8+NKG2A cells (ρ = 0.89, p = 0.0333). The right panel (**B**) shows the negative correlation between baseline Shannon diversity and percent change in frequency of vRNA^+^ cells of rectum (ρ = −0.67, p = 0.0394). Each point represents an individual sample, with trend lines and 95% confidence intervals. Spearman correlation coefficients (ρ) and p-values are displayed for each relationship.

To explore possible changes in specific microbial features following N-803 treatment, we applied *MaAsLin2* mixed-effects modeling to 413 taxonomic profiles against 10 immune markers and the frequency of tissue-associated vRNA^+^ cells. At FDR *q* < 0.25, we identified 36 taxa associated with vRNA^+^ cell frequency (see Supplementary Materials Table S3) in all 3 tissues and 90 taxa linked to immune parameters (see Supplementary Materials Table S4). Taxa such as *Faecalibacterium* HTFF (q = 0.2), *Lachnospiraceae* (q = 0.2), *Roseburia* spp (q = 0.11), *Anaerotignum faecicola* (q = 0.15), *Clostridiales* (q = 0.16), and *Oscillospiraceae* (q = 0.02) demonstrated strong positive associations with the cytotoxic marker Perforin and activation markers CD69 and Ki-67^+^ on CD8^+^ T cells (Figure 6A). Conversely, *Prevotella dentalis* (q = 0.13), *Dialister invisus* (q = 0.08), and *Lawsonibacter assacharolyticus* (q = 0.1) were negatively correlated with NK cell FasL expression and CD8^+^ T cell Ki-67 expression (Figure 6A). In terms of viral persistence, taxa such as *Lawsonibacter asaccharolyticus* (q = 0.06) and *Megasphaera* spp (q = 0.0002) exhibited positive correlations with the frequency of vRNA^+^ cells in rectal and ileal tissues (Figure 6B). Further modeling of metabolic pathways identified several SCFA-related fermentation pathways positively associated with markers of cytotoxic immune-cell activation (CD8^+^CD69^+^) and proliferation (CD8^+^Ki-67^+^), such as PWY 5494 (pyruvate fermentation to propanoate; q = 0.18), PROPFERM PWY (Stickland reaction; q = 0.17; Fig. 6C; Supplementary Materials Table S5). Interestingly, SCFA-related pathways were negatively associated with vRNA^+^ cell frequency in the tissues (*q* < 0.25, Supplementary Materials Table S6). Collectively, these data suggest that baseline microbiome diversity, as well as specific SCFA-producing taxa and fermentation pathways, are associated with both increased immune activation markers and reduced frequency of tissue vRNA^+^ cells following N-803 treatment.

**Figure 6:**
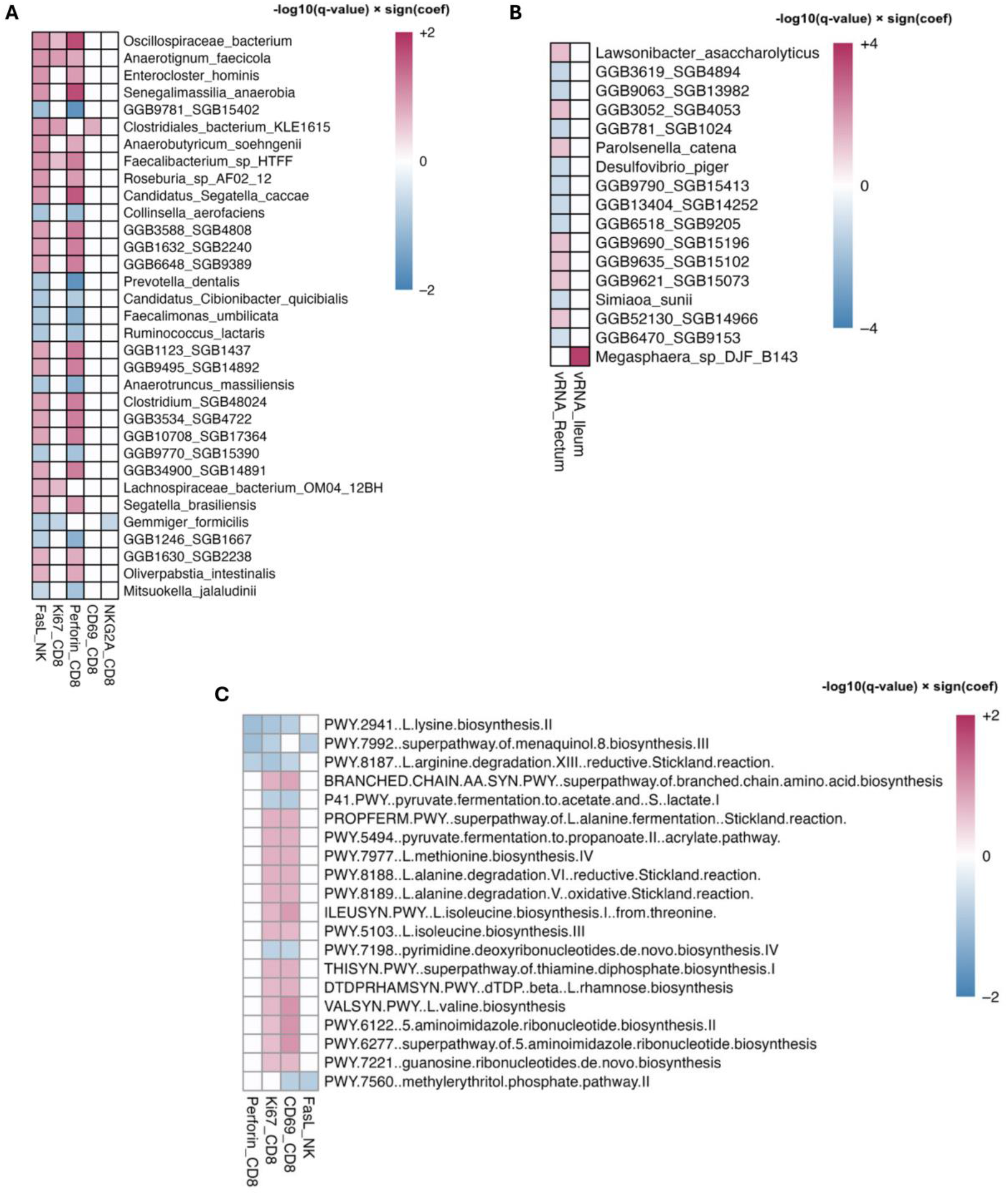
Differentially abundant bacterial species and microbial pathways obtained from MaAsLin2 analysis. The bacterial species correlated with multiple metadata fixed effects. The red color shows a positive association, and the blue color shows a negative association. **(A)** Heatmap displaying the species significantly associated (*q* < *0*.*25*) with the functional markers of CD8 and NK cells. **(B)** Heatmap displaying the species significantly associated (*q* < *0*.*1*) with the frequency of vRNA^+^ cells in the rectum and ileum. **(C)** Heatmap displaying the microbial pathways significantly associated (*q* < *0*.*25*) with the functional markers of CD8 and NK cells.

## Discussion

Our longitudinal study shows that three subcutaneous doses of the IL-15 superagonist N-803 did not significantly modify the overall structure or function of the fecal microbiome in ART-suppressed PWH. These findings are in agreement with previous COVID-19 vaccine studies and cancer immunotherapy research, where bacterial diversity and composition were largely unaltered following the intervention [20,30-32]. This is also consistent with the clinical outcomes from the Phase 1B trial, where N-803 was well tolerated by the participants and resulted in no reported gastrointestinal adverse events [18], further supporting the mucosal-enhancing safety profile of N-803.

While community-level diversity metrics were mostly stable throughout the study, *MaAsLin2* modeling was able to identify subtle shifts in specific taxa and metabolic pathways. The relative abundance of *Faecalibacterium prausnitzii*, a key butyrate-producing bacterium known for anti-inflammatory and barrier-supportive roles [24], was significantly elevated following N-803 administration. This enrichment may signal a shift toward a more favorable mucosal environment, potentially supported by the significant decline in vRNA^+^ cell frequency observed in both gut and lymph node tissues in the Phase 1B trial [18]. PWH often experience a depletion of *F. prausnitzii* that persists even during suppressive ART and has been linked to reduced SCFA production and compromised gut mucosal integrity [33,34]. Although the SCFA fermentation pathways showed only nominal enrichment trends in our study, these trends paralleled the observed expansion of *F. prausnitzii*, reflecting a coordinated functional adaptation. These microbial metabolites are key to keeping the epithelial barrier intact, regulating mucosal immunity [25] and HIV-associated comorbidities [35].

Conversely, the relative abundance of *Segatella sinensis* decreased after N-803. The members of the *Segatella/Prevotella* clade have been implicated in mucosal inflammation and increased gut permeability in PWH [36]. Cancer immunotherapy studies have reported mixed results between *Prevotella* abundance and treatment outcome. For instance, an early melanoma study showed *Prevotella*-rich communities linked to anti-PD-1 therapy non-responders [19], while studies by Peng *et al*. and Pietrzak *et al*. have reported higher *Prevotella* abundance in anti-PD-1 therapy responders [20,37]. In this study, the decrease in *S. sinensis* may indicate a shift toward a less inflammatory gut environment. Given the limited literature on *S. sinensis*, further investigation is necessary to determine whether its reduction directly enhances mucosal immunity or reflects N-803-mediated effects.

Likewise, N-803 administration led to a significant reduction in the abundance of the microbial histidine degradation pathway, suggesting a shift towards a less inflammatory gut environment in ART-suppressed PWH. Previous research has shown that elevated microbial histidine metabolism drives systemic inflammation, primarily through histamine-driven inflammasome modulation [38] and pro-inflammatory metabolite release, such as imidazole propionate [39]. The concurrent decrease in this pro-inflammatory pathway, alongside trends in SCFA pathway enrichment, suggests that N-803 promotes a functional rebalancing of the gut microbiome toward a more immunologically favorable state. The shift observed in these pathways, despite a largely unaltered microbiome composition, suggests that N-803-mediated immunologic effects selectively support the recovery or expansion of taxa and microbial functions essential for gut homeostasis.

The central finding of our study is that higher baseline gut microbial diversity correlated with both immunological activation and reduced frequency of tissue vRNA+ cells following N-803 therapy. Participants with higher alpha diversity exhibited increased CD8^+^ T-cell activation (higher CD69 and NKG2A) and enhanced NK-cell cytotoxic function (elevated FasL). Furthermore, baseline diversity inversely correlated with the frequency of vRNA^+^ cells in rectal tissues post-treatment, suggesting that a more diverse microbiome may facilitate more effective immune-mediated viral suppression. To our knowledge, this is the first N-803 study in an HIV cohort demonstrating a correlation between baseline gut microbiome diversity and its metabolic functions with treatment outcomes. This finding is in line with previous studies in cancer therapy and transplant, where multiple studies have shown that higher baseline microbiome diversity correlated with positive responses to treatment [19–21,37,40–42]. The concept of pre-treatment diversity as a predictive biomarker also transcends cancer therapy: multiple vaccine studies have shown that individuals with greater pre-vaccination microbial diversity and higher levels of SCFAs exhibit better vaccine-induced immune responses [30,31,43,44].

Interestingly, many of the same taxa and pathways that shifted longitudinally after N-803 treatment also correlated with post-treatment immune functional markers. For instance, SCFA-producing bacterial taxa, such as *Faecalibacterium* spp., *Roseburia* spp., *Lachnospiraceae*, and *Oscillospiraceae*, were positively associated with functional markers of activated CD8^+^ and NK cells. These taxa have previously been implicated in positive immunotherapy outcomes [21]. In a colorectal cancer study, *Roseburia intestinalis* was shown to enhance CD8^+^ T cell function and improve responses to checkpoint blockade [45]. The negative association between the abundance of *Lawsonibacter asaccharolyticus* and FasL expression, coupled with its positive association with the frequency of vRNA^+^ cells in rectal tissues, suggests this species could be a biomarker for weakened immune responses. This is supported by a vaccination study where *L. asaccharolyticus* was enriched in low responders and negatively correlated with neutralizing antibody production [30]. Microbial pathways related to SCFA biosynthesis (e.g., pyruvate fermentation to propanoate, Stickland reactions) were also enriched after N-803 treatment and positively associated with immune activation markers. These pathway-level findings reinforce the idea that microbiome function, especially SCFA metabolism, could be a key factor influencing the host response to immunotherapy [20,25,43].

Our data suggest that individuals with greater microbial diversity may be primed to respond more effectively to N-803 therapy. N-803 administration subtly enriches a subset of immunologically beneficial microbes and pathways without destabilizing the overall community. However, these findings must be considered in the context of our study’s limitations. Our cohort was small and predominantly male, limiting its generalizability and lacking the power to detect longitudinal patterns with greater statistical confidence. We lacked detailed dietary information and metabolomic profiling, and relied on stool sampling, which may have resulted in missing important mucosal changes. Future studies should include larger cohorts, integrate mucosal metagenomics data, and employ mechanistic models to clarify causal mechanisms underlying microbiome-immune interactions and further explore microbiome modulation to enhance therapeutic efficacy.

## Conclusions

In conclusion, our data adds to the growing body of research showing the gut microbiome as a key factor in determining immunotherapy success in PWH. Baseline microbial diversity and specific microbial pathways seem to influence the level of immune activation and virus transcriptional control after N-803. These results indicate that the gut microbiome is not a passive indicator of treatment progression but may actively impact immunotherapeutic outcomes. Importantly, they suggest that improving microbiome diversity or function before N-803 treatment could boost immunologic and virologic responses in ART-suppressed PWH. These findings provide a basis for microbiome-based strategies to enhance HIV cure efforts through tailored immunotherapy.

## Acknowledgments

We thank all the participants who agreed to participate and provide samples for this study. Shotgun sequencing was performed by the University of Minnesota Genomics Center (UMGC). We also acknowledge the Minnesota Supercomputing Institute (MSI) at the University of Minnesota for providing computational resources that contributed to the research results reported within this paper.

## Data availability

Raw shotgun metagenomics sequencing data is available at NCBI under BioProject ID PRJNA1289810. All code and data used for this analysis are available upon request.

## Author Contributions

Conceptualization: NRK, TWS

Supervision: NRK, TWS

Lab support: NRK, TWS, JSM

Lab Investigation: CMB, TAS, RC, JA, KE, JR

Data analysis: AC

Visualization: AC, RSC

Writing-original draft: AC

Writing-review & editing: AC, RC, CMB, JA, TAS, KE, RSC, RL, ES, JSM, JR, TWS, NRK

## References

[1] Estes JD, Kityo C, Ssali F, et al. Defining total-body AIDS-virus burden with implications for curative strategies. Nat Med 2017;23:1271–6. 10.1038/nm.4411.

[2] Buzon MJ, Martin-Gayo E, Pereyra F, et al. Long-term antiretroviral treatment initiated at primary HIV-1 infection affects the size, composition, and decay kinetics of the reservoir of HIV-1-infected CD4 T cells. J Virol 2014;88:10056–65. 10.1128/JVI.01046-14.

[3] Gantner P, Buranapraditkun S, Pagliuzza A, et al. HIV rapidly targets a diverse pool of CD4+ T cells to establish productive and latent infections. Immunity 2023;56:653–668.e5. 10.1016/j.immuni.2023.01.030.

[4] Fletcher CV, Staskus K, Wietgrefe SW, et al. Persistent HIV-1 replication is associated with lower antiretroviral drug concentrations in lymphatic tissues. Proc Natl Acad Sci U S A 2014;111:2307–12. 10.1073/pnas.1318249111.

[5] Lorenzo-Redondo R, Fryer HR, Bedford T, et al. Persistent HIV-1 replication maintains the tissue reservoir during therapy. Nature 2016;530:51–6. 10.1038/nature16933.

[6] Prigann J, Tavora R, Furler O’Brien RL, et al. Silencing the transcriptionally active HIV reservoir to improve treatment outcomes. Nat Microbiol 2024;9:2470–2. 10.1038/s41564-024-01816-5.

[7] Cohen GB, Gandhi RT, Davis DM, et al. The Selective Downregulation of Class I Major Histocompatibility Complex Proteins by HIV-1 Protects HIV-Infected Cells from NK Cells. Immunity 1999;10:661–71. 10.1016/S1074-7613(00)80065-5.

[8] Körner C, Granoff ME, Amero MA, et al. Increased frequency and function of KIR2DL1-3+ NK cells in primary HIV-1 infection are determined by HLA-C group haplotypes. Eur J Immunol 2014;44:2938–48. 10.1002/eji.201444751.

[9] Körner C, Simoneau CR, Schommers P, et al. HIV-1-Mediated Downmodulation of HLA-C Impacts Target Cell Recognition and Antiviral Activity of NK Cells. Cell Host Microbe 2017;22:111–119.e4. 10.1016/j.chom.2017.06.008.

[10] Verbist KC, Klonowski KD. Functions of IL-15 in anti-viral immunity: Multiplicity and variety. Cytokine 2012;59:467–78. 10.1016/j.cyto.2012.05.020.

[11] Guo Y, Luan L, Patil NK, et al. Immunobiology of the IL-15/IL-15Rα complex as an antitumor and antiviral agent. Cytokine Growth Factor Rev 2017;38:10–21. 10.1016/j.cytogfr.2017.08.002.

[12] Harwood O, O’Connor S. Therapeutic Potential of IL-15 and N-803 in HIV/SIV Infection. Viruses 2021;13:1750. 10.3390/v13091750.

[13] Miller JS, Davis ZB, Helgeson E, et al. Safety and virologic impact of the IL-15 superagonist N-803 in people living with HIV: a phase 1 trial. Nat Med 2022;28:392–400. 10.1038/s41591-021-01651-9.

[14] McBrien JB, Mavigner M, Franchitti L, et al. Robust and persistent reactivation of SIV and HIV by N-803 and depletion of CD8+ cells. Nature 2020;578:154–9. 10.1038/s41586-020-1946-0.

[15] Miller JS, Rhein J, Davis ZB, et al. Safety and Virologic Impact of Haploidentical NK Cells Plus Interleukin 2 or N-803 in HIV Infection. J Infect Dis 2024;229:1256–65. 10.1093/infdis/jiad578.

[16] Jones RB, Mueller S, O’Connor R, et al. A Subset of Latency-Reversing Agents Expose HIV-Infected Resting CD4+ T-Cells to Recognition by Cytotoxic T-Lymphocytes. PLOS Pathog 2016;12:e1005545. 10.1371/journal.ppat.1005545.

[17] Ellis-Connell AL, Balgeman AJ, Zarbock KR, et al. ALT-803 Transiently Reduces Simian Immunodeficiency Virus Replication in the Absence of Antiretroviral Treatment. J Virol 2018;92:10.1128/jvi.01748-17. 10.1128/jvi.01748-17.

[18] Rhein J, Chipman JG, Beilman GJ, et al. Impact of the IL-15 superagonist N-803 on lymphatic reservoirs of HIV. JCI Insight 2025. 10.1172/jci.insight.190831.

[19] Gopalakrishnan V, Spencer CN, Nezi L, et al. Gut microbiome modulates response to anti-PD-1 immunotherapy in melanoma patients. Science 2018;359:97–103. 10.1126/science.aan4236.

[20] Peng Z, Cheng S, Kou Y, et al. The Gut Microbiome Is Associated with Clinical Response to Anti-PD-1/PD-L1 Immunotherapy in Gastrointestinal Cancer. Cancer Immunol Res 2020;8:1251–61. 10.1158/2326-6066.CIR-19-1014.

[21] Zhang M, Liu J, Xia Q. Role of gut microbiome in cancer immunotherapy: from predictive biomarker to therapeutic target. Exp Hematol Oncol 2023;12:84. 10.1186/s40164-023-00442-x.

[22] Fluckiger A, Daillère R, Sassi M, et al. Cross-reactivity between tumor MHC class I-restricted antigens and an enterococcal bacteriophage. Science 2020;369:936–42. 10.1126/science.aax0701.

[23] Zhou J, Zhang Y, Cui P, et al. Gut Microbiome Changes Associated With HIV Infection and Sexual Orientation. Front Cell Infect Microbiol 2020;10. 10.3389/fcimb.2020.00434.

[24] Sokol H, Pigneur B, Watterlot L, et al. Faecalibacterium prausnitzii is an anti-inflammatory commensal bacterium identified by gut microbiota analysis of Crohn disease patients. Proc Natl Acad Sci 2008;105:16731–6. 10.1073/pnas.0804812105.

[25] Ney L-M, Wipplinger M, Grossmann M, et al. Short chain fatty acids: key regulators of the local and systemic immune response in inflammatory diseases and infections. Open Biol 2023;13:230014. 10.1098/rsob.230014.

[26] Klatt NR, Funderburg NT, Brenchley JM. Microbial translocation, immune activation, and HIV disease. Trends Microbiol 2013;21:6–13. 10.1016/j.tim.2012.09.001.

[27] Swanson EC, Basting CM, Klatt NR. The role of pharmacomicrobiomics in HIV prevention, treatment, and women’s health. Microbiome 2024;12:254. 10.1186/s40168-024-01953-3.

[28] Vujkovic-Cvijin I, Somsouk M. HIV and the Gut Microbiota: Composition, Consequences, and Avenues for Amelioration. Curr HIV/AIDS Rep 2019;16:204–13. 10.1007/s11904-019-00441-w.

[29] Langat R, Chakrawarti A, Klatt NR. Cannabis Use in HIV: Impact on Inflammation, Immunity and the Microbiome. Curr HIV/AIDS Rep 2025;22:19. 10.1007/s11904-025-00729-0.

[30] Tang B, Tang L, He W, et al. Correlation of gut microbiota and metabolic functions with the antibody response to the BBIBP-CorV vaccine. Cell Rep Med 2022;3:100752. 10.1016/j.xcrm.2022.100752.

[31] Boston RH, Guan R, Kalmar L, et al. Stability of gut microbiome after COVID-19 vaccination in healthy and immuno-compromised individuals. Life Sci Alliance 2024;7:e202302529. 10.26508/lsa.202302529.

[32] Zhu C, Zhang C, Wang S, et al. Characterizations of multi-kingdom gut microbiota in immune checkpoint inhibitor-treated hepatocellular carcinoma. J Immunother Cancer 2024;12. 10.1136/jitc-2023-008686.

[33] Dubourg G, Lagier J-C, Hüe S, et al. Gut microbiota associated with HIV infection is significantly enriched in bacteria tolerant to oxygen. BMJ Open Gastroenterol 2016;3:e000080. 10.1136/bmjgast-2016-000080.

[34] Serrano-Villar S, Vázquez-Castellanos JF, Vallejo A, et al. The effects of prebiotics on microbial dysbiosis, butyrate production and immunity in HIV-infected subjects. Mucosal Immunol 2017;10:1279–93. 10.1038/mi.2016.122.

[35] Sereti I, Verburgh ML, Gifford J, et al. Impaired gut microbiota-mediated short-chain fatty acid production precedes morbidity and mortality in people with HIV. Cell Rep 2023;42. 10.1016/j.celrep.2023.113336.

[36] Vázquez-Castellanos JF, Serrano-Villar S, Latorre A, et al. Altered metabolism of gut microbiota contributes to chronic immune activation in HIV-infected individuals. Mucosal Immunol 2015;8:760–72. 10.1038/mi.2014.107.

[37] Pietrzak B, Tomela K, Olejnik-Schmidt A, et al. A Clinical Outcome of the Anti-PD-1 Therapy of Melanoma in Polish Patients Is Mediated by Population-Specific Gut Microbiome Composition. Cancers 2022;14:5369. 10.3390/cancers14215369.

[38] Levy M, Thaiss CA, Zeevi D, et al. Microbiota-Modulated Metabolites Shape the Intestinal Microenvironment by Regulating NLRP6 Inflammasome Signaling. Cell 2015;163:1428–43. 10.1016/j.cell.2015.10.048.

[39] Koh A, Molinaro A, Ståhlman M, et al. Microbially Produced Imidazole Propionate Impairs Insulin Signaling through mTORC1. Cell 2018;175:947–961.e17. 10.1016/j.cell.2018.09.055.

[40] Otto-Dobos LD, Strehle LD, Loman BR, et al. Baseline gut microbiome alpha diversity predicts chemotherapy-induced gastrointestinal symptoms in patients with breast cancer. Npj Breast Cancer 2024;10:1–12. 10.1038/s41523-024-00707-6.

[41] Sims TT, El Alam MB, Karpinets TV, et al. Gut microbiome diversity is an independent predictor of survival in cervical cancer patients receiving chemoradiation. Commun Biol 2021;4:1–10. 10.1038/s42003-021-01741-x.

[42] Taur Y, Jenq RR, Perales M-A, et al. The effects of intestinal tract bacterial diversity on mortality following allogeneic hematopoietic stem cell transplantation. Blood 2014;124:1174–82. 10.1182/blood-2014-02-554725.

[43] Daddi L, Dorsett Y, Geng T, et al. Baseline Gut Microbiome Signatures Correlate with Immunogenicity of SARS-CoV-2 mRNA Vaccines. Int J Mol Sci 2023;24:11703. 10.3390/ijms241411703.

[44] Wilson A, Manuzak JA, Liang H, et al. Probiotic Therapy During Vaccination Alters Antibody Response to Simian-Human Immunodeficiency Virus Infection But Not to Commensals. AIDS Res Hum Retroviruses 2023;39:222–31. 10.1089/aid.2022.0123.

[45] Kang X, Liu C, Ding Y, et al. Roseburia intestinalis generated butyrate boosts anti-PD-1 efficacy in colorectal cancer by activating cytotoxic CD8+ T cells. Gut 2023;72:2112. 10.1136/gutjnl-2023-330291.

