## Supplementary Materials for "Pre-treatment Microbiome Diversity and Function is associated with Expansion of Cytotoxic and Regulatory Immune Populations after N-803 treatment in People with HIV"

**Supplementary Methods**

**Fecal Sample Collection**

Fecal samples for microbiome analysis were longitudinally collected from participants at defined intervals throughout N-803 administration (Figure 1). Samples were collected at Baseline (before the initial dose of N-803), following the second dose (Dose2), immediately after the completion of all three doses (Postdrug), and at a follow-up visit approximately three months post-treatment (Month3). Samples were collected using the OMNIgene®•GUT collection kit (OMR-200, DNA Genotek, Ottawa, Canada) and stored at 80°C until DNA extraction was performed, using the Qiagen PowerSoil Pro extraction kit (Qiagen, Germantown, USA).

**Shotgun Metagenomics Analysis of Gut Microbiota**

Total genomic DNA was extracted from fecal samples using the DNeasy 96 PowerSoil Pro QIAcube HT kit (Qiagen, Germantown, USA) on a QIAcube following the manufacturer's instructions. Shotgun metagenomic sequencing was performed by the University of Minnesota Genomics Core (UMGC). Briefly, the libraries were prepared using the Illumina NexteraXT DNA library preparation kit (Cat. #FC-131-1096) according to the manufacturer's guidelines. Sequencing was performed on an Illumina NovaSeq SP platform to generate 150 bp paired-end reads from 40 dual-indexed NexteraXT libraries. A total of 872 million paired-end reads were generated across all samples.

The quality of raw sequencing data was assessed using FastQC (v0.11.9). Adapter sequences and low-quality bases (Phred score <30) were removed through TrimGalore (v0.6.10). High-quality reads were subsequently aligned to the reference human genome GRCh38 to filter out host reads using Bowtie2 (v4.2.5), and the resulting alignments were further processed using SAMtools (v1.17).

Taxonomic profiling of microbial communities was conducted using two parallel approaches: we used Kraken2 (v2.1.3) with a confidence score threshold of 0.2, followed by relative abundance estimation using Bracken (v3.0.1), and also MetaPhlAn4. Beta diversity patterns across timepoints were assessed using Bray–Curtis distances, and individual-level trajectories were visualized for both methods. Temporal trends in compositional shifts were consistent between Kraken2 and MetaPhlAn4 (Supplementary Figure S1 A-D). To formally test for concordance between methods, we performed a Procrustes analysis on the first two principal coordinate axes of each dataset. The overlay of corresponding samples revealed a strong concordance between Kraken2 and MetaPhlAn4 ordinations (Procrustes correlation = 0.521, *p* = 0.001) (Figure S1A), indicating microbiome compositional structures recovered by the two methods were more similar than would be expected by chance. Based on this agreement between two taxonomic classifiers, and the advantage of unified databases and methods for downstream functional analyses, MetaPhlAn4-derived taxonomic profiles were utilized for all subsequent analyses reported here. Functional pathways and gene families were profiled using HUMAnN3 (v3.9), yielding MetaCyc pathway abundances. Species-level taxonomic profiles obtained from MetaPhlAn4 that were present in fewer than 5% of samples or with a relative abundance below 0.05% were removed to reduce spurious observations. Both taxonomic and functional abundance tables were imported into *phyloseq* (v1.46.0), and relative abundance and centered log-ratio (CLR) transformations were applied using *microbiome* (v.24.0).

**Statistical Analysis**

Alpha diversity was quantified using Shannon and Simpson indices, calculated on both taxonomic and functional abundances, and temporal trends were assessed using *lme4* (v 1.1.35.3) linear mixed‐effects models with participants as a random intercept. Overall effects of sampling time (Baseline, Dose2, Postdrug, Month3) were evaluated by ANOVA on the fitted models and pairwise contrasts between timepoints were extracted using *emmeans* (v1.10.2) with Benjamini–Hochberg (BH) adjustment for multiple comparisons. Beta diversity was calculated using Aitchison distances on CLR-transformed abundance tables and Bray-Curtis dissimilarities on proportional abundance data, with *vegan* (v2.6.4). Ordination was visualized via Principal Coordinates Analysis (PCoA), and pairwise PERMANOVA tests were performed to identify significant compositional shifts between timepoints using *pairwiseAdonis* (0.4.1), with resulting p-values BH-adjusted. Differential abundance analysis was performed with *MaAsLin2* (v1.16.0), applying mixed-effects modeling with participant as a random effect and *p*-values BH adjusted.

To assess the relationship between baseline microbiome diversity and post N-803 immune responses, we performed Spearman correlation analyses between Baseline Shannon diversity and i) the Postdrug expression levels of immune markers on NK cells and CD8+ T cells, and ii) the percent change in marker expression from Baseline to Postdrug for each participant. These included markers associated with cytotoxicity (Perforin, Granzyme B), activation (CD38, HLA-DR), proliferation (Ki-67), differentiation and senescence (CD27, CD57), inhibitory signaling (PD-1, NKG2A), and apoptosis (FasL). CyTOF-derived expression values were treated as continuous variables and reported as the mean marker intensity (MMI) within the respective immune cell subset. Correlations were computed using *psych* (v2.4.6.26), with nominal p-values <0.05 considered significant.

The BH-adjusted q-values ≤ 0.25 were considered significant for all multiple-testing analyses due to the small sample size. All analyses were conducted in R (v4.3.3), and data visualizations were generated using *ggplot2* (v3.5.1) and *pheatmap* (v1.0.12).

**Supplementary** **figures**

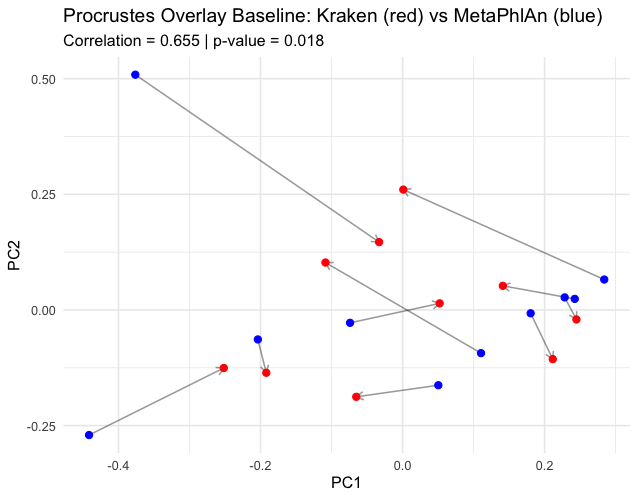

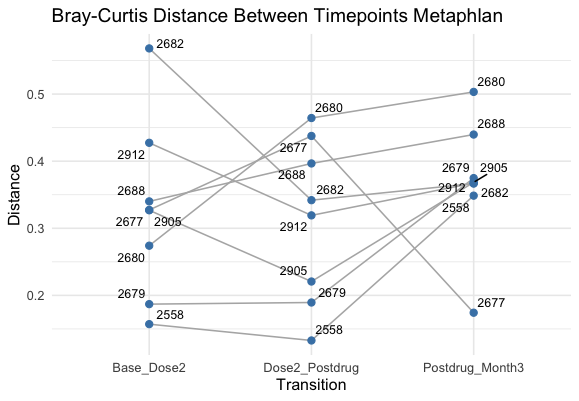

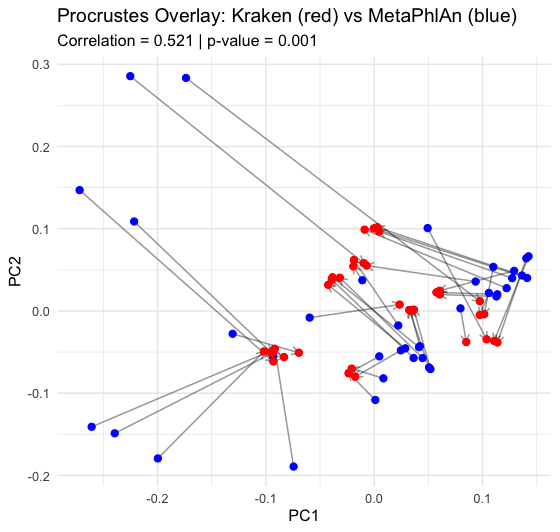

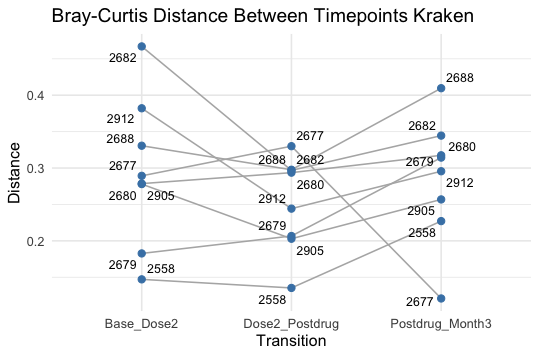

A

C

D

B

**Figure S1: Concordance between Kraken2 and MetaPhlAn 4 taxonomic ordinations and longitudinal compositional shifts. (A)** Procrustes overlay of Bray–Curtis PCoA ordinations for all 38 stool samples. Kraken2 points are shown in red, MetaPhlAn 4 in blue; grey line segments connect the two profiles derived from the same individual. The symmetric Procrustes correlation was 0.521 (999 permutations, p = 0.001). **(B)** The same Procrustes analysis was restricted to Baseline samples (ρ = 0.655, p = 0.018). **(C)** Within-participant Bray–Curtis distances between successive time-points computed from Kraken2 data; points are labelled with participant IDs and joined by grey lines to illustrate individual trajectories. **(D)** Equivalent distance plot using MetaPhlAn 4 profiles. Similar spread and ordering of distances in panels C and D underscore the consistency of longitudinal patterns recovered by the two approaches.

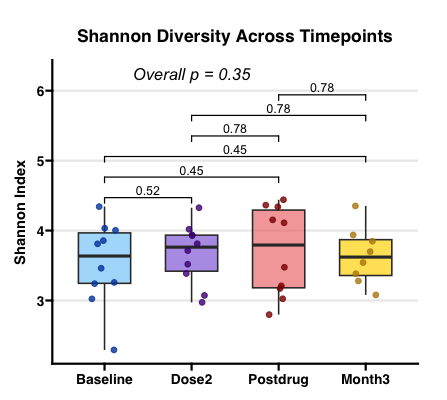

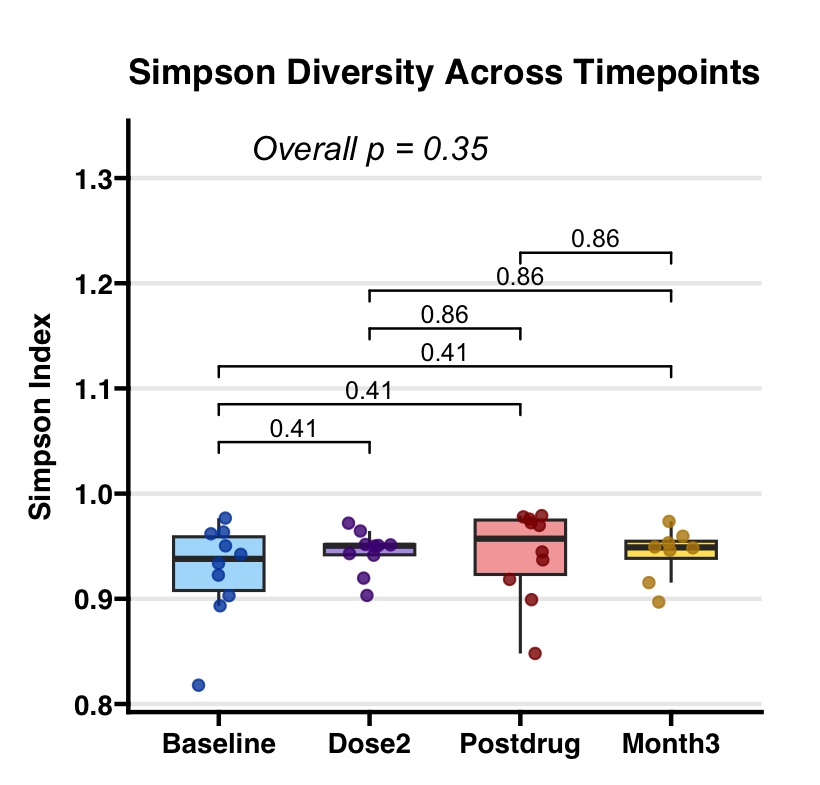

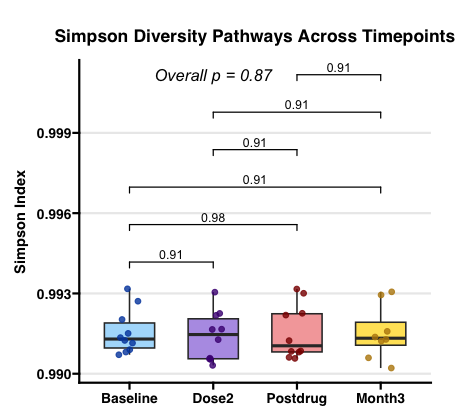

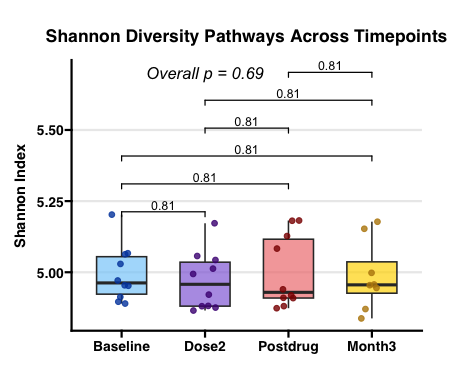

**A**

**C**

**B**

**D**

**Figure S2: Longitudinal α-diversity of the fecal microbiome and its metabolic pathways during N-803 therapy.** **(A)** Shannon and **(B)** Simpson species-level diversity derived from MetaPhlAn4 profiles remain stable from Baseline through Month 3 (linear mixed-effects model, overall *p* = 0.35 for both metrics). No pairwise Wilcoxon comparison between consecutive visits reaches significance after BH adjustment. **(C)** Simpson and **(D)** Shannon pathway-level diversity calculated from HUMAnN3 MetaCyc profiles, likewise show no significant temporal change (overall *p* = 0.87 and *p* = 0.69, respectively). Boxes indicate the inter-quartile range with the median as a horizontal bar; whiskers extend to 1.5 × IQR; points represent individual participants.

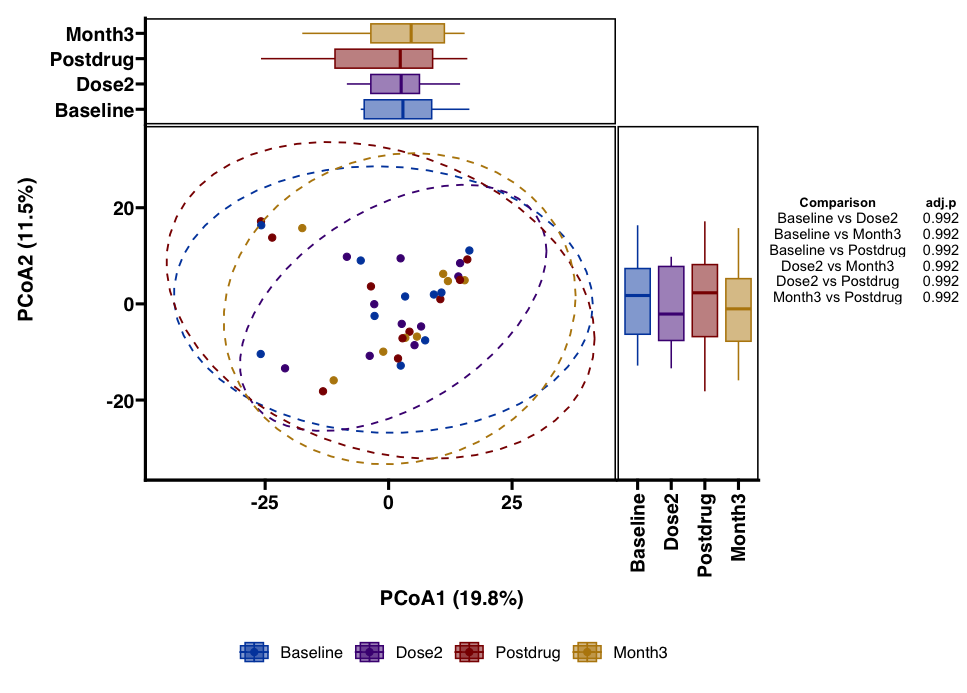

**Figure S3:** Principal Coordinates Analysis (PCoA) plot based on Euclidean distances of CLR transformed MetaCyc pathway abundances. Each point represents a sample, colored by timepoint, with 95% confidence ellipses. Side boxplots show the distribution of PCoA1 and PCoA2 scores by timepoint. Pairwise PERMANOVA results are summarized with adjusted p-values (FDR) for each timepoint comparison.

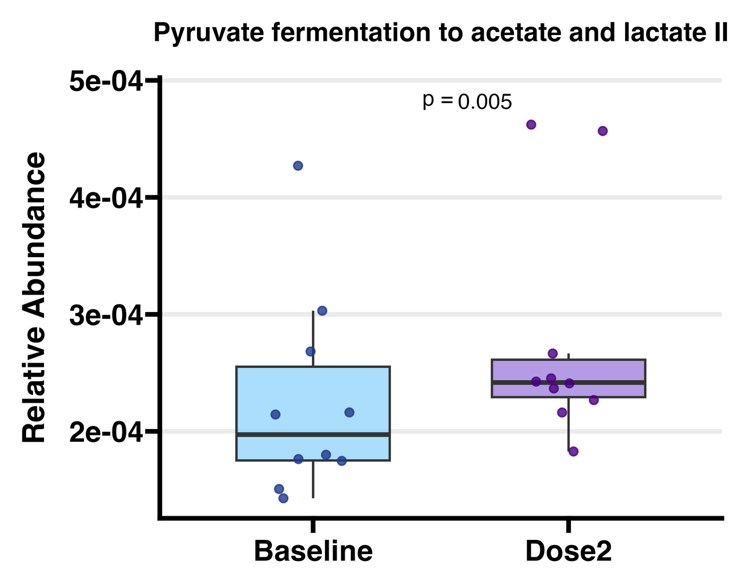

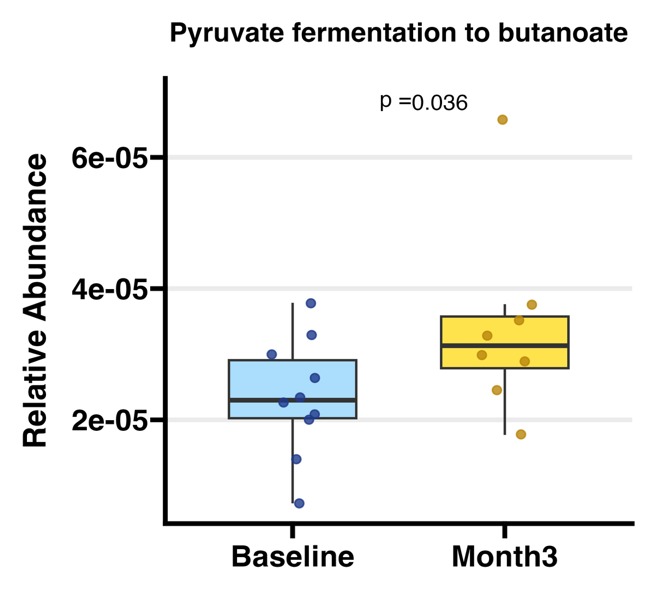

**Figure S4: Differential abundance of fecal bacterial MetaCyc pathways after N-803 therapy.** Box-and-whisker plots show relative abundances (HUMAnN 3). Short-chain fatty acid pathways increased after treatment (nominal p < 0.05): pyruvate-to-acetate/lactate increased by ~15 % from Baseline to Dose 2, and pyruvate-to-butanoate increased by ~45 % from Baseline to Month 3.

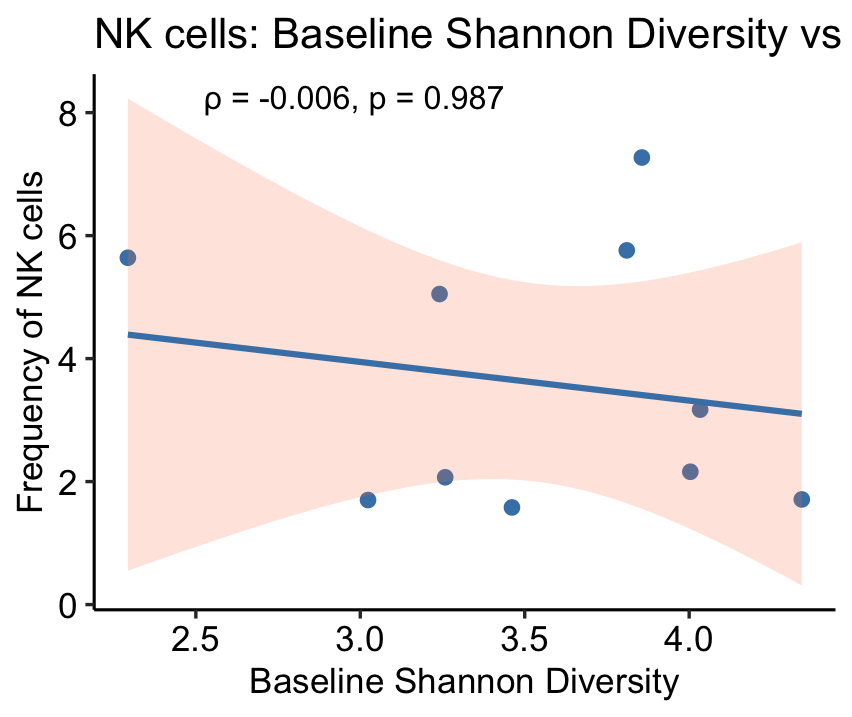

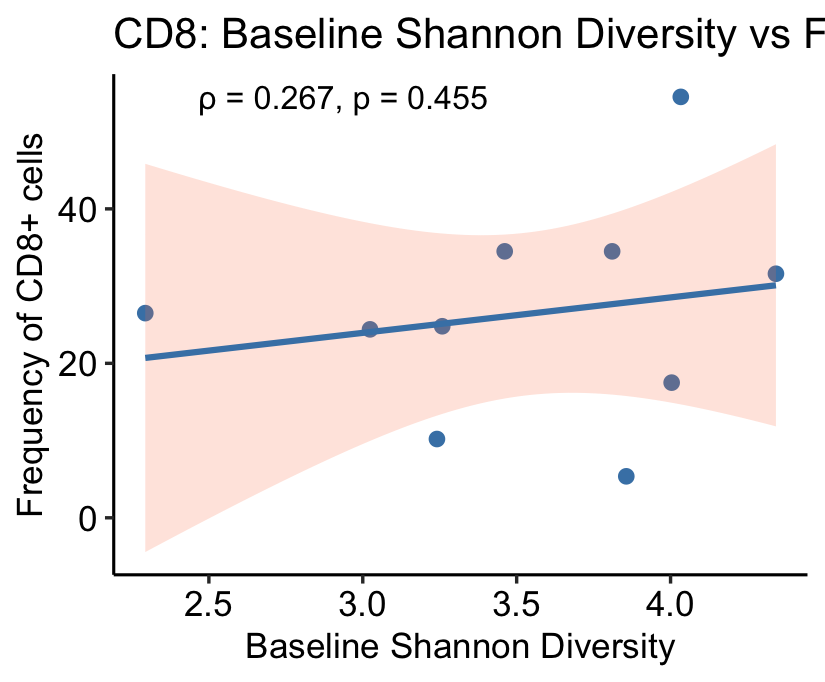

**A**

**B**

**Figure S5: Baseline gut Shannon diversity versus lymph-node immune composition.** Scatter plots relate baseline fecal Shannon diversity (x-axis) to the Postdrug percentage in frequency of CD8 ⁺ T cells **(A)** or NK cells **(B)** within lymph-node biopsy–derived mononuclear cells (LNBCs). Each point represents an individual participant; solid blue lines depict least-squares fits with 95 % confidence bands. A weak positive trend is evident for CD8⁺ T-cell frequency (Spearman ρ = 0.267, p = 0.455), whereas NK-cell frequency shows no association (ρ = -0.006, p = 0.987).

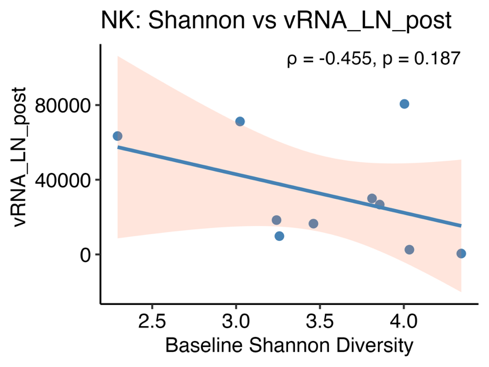

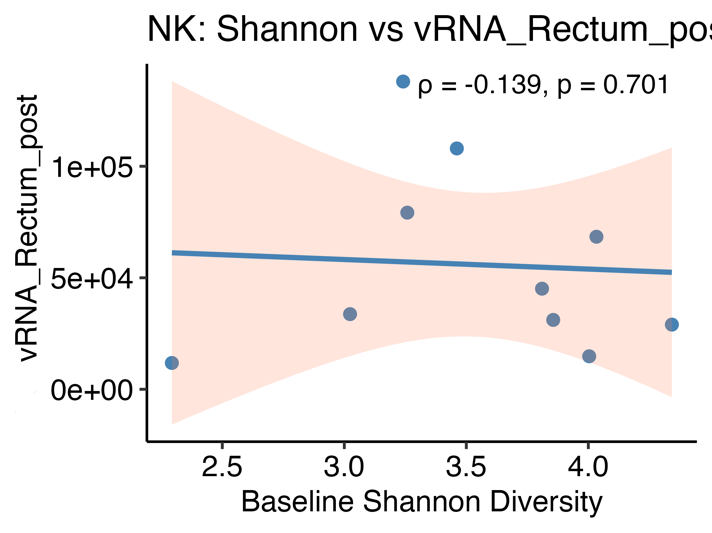

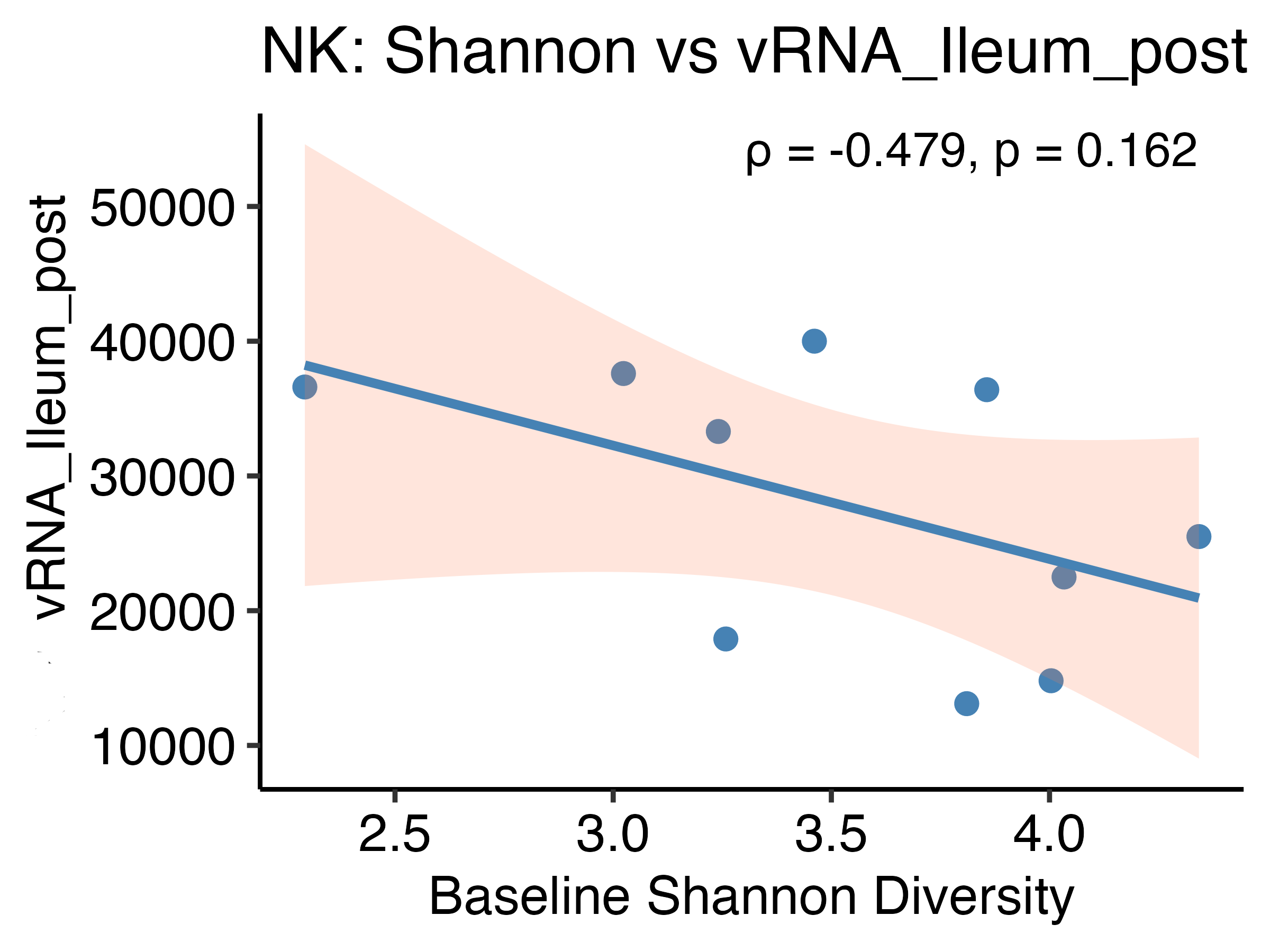

**A**

**C**

**D**

**E**

**B**

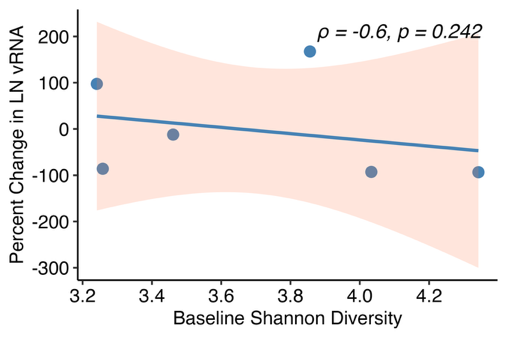

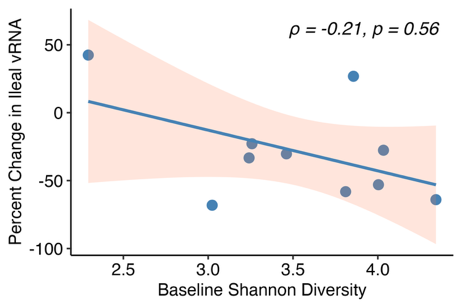

**Figure S6: Baseline fecal Shannon diversity and tissue‐associated frequency of HIV RNA+ cells after N-803 therapy.** Scatter plots correlate baseline fecal Shannon diversity (x-axis) with the frequency of HIV RNA+ cells in lymph-node (LN), ileal, and rectal CD4⁺ T-cell compartments. Panels A-B show the **percent change** in frequency of HIV RNA+ cells from Baseline to the Postdrug biopsy (LN: ρ = -0.60, p = 0.242; ileum: ρ = -0.21, p = 0.560; n = 6-8). Panels C-E display the **absolute vRNA levels per sample** at the Postdrug visit (LN: ρ = -0.455, p = 0.187; rectum: ρ = -0.139, p = 0.701; ileum: ρ = -0.479, p = 0.162; n = 8). Points represent individual participants; solid blue lines denote least-squares fits with shaded 95 % confidence bands. All correlations are negative, but none reach statistical significance.
